# Logical modelling of myelofibrotic microenvironment predicts dysregulated progenitor stem cell crosstalk

**DOI:** 10.1101/2022.12.12.520119

**Authors:** S. P. Chapman, E. Duprez, E. Remy

## Abstract

Primary myelofibrosis is an untreatable age-related disorder of haematopoiesis in which a break in the crosstalk between progenitor Haematopoietic Stem Cells (HSCs) and neighbouring mesenchymal stem cells causes HSCs to rapidly proliferate and migrate out of the bone marrow. 90% of patients harbour mutations in driver genes that all converge to overactivate hematopoietic JAK-STAT signalling, which is thought to be critical for disease progression, as well as microenvironment modification induced by chronic inflammation. The trigger to the initial event is unknown but dysregulated thrombopoietin (TPO) and Toll-Like Receptor (TLR) signalling are hypothesised to initiate chronic inflammation which then disrupts stem cell crosstalk. Using a systems biology approach, we have constructed an inter and intracellular logical model that captures JAK-STAT signalling and key crosstalk channels between haematopoietic and mesenchymal stem cells. The aim of the model is to decipher how TPO and TLR stimulation can perturb the bone marrow microenvironment and dysregulate stem cell crosstalk. The model predicted conditions in which the disease was averted and established for both wildtype and ectopically JAK mutated simulations. The presence of TPO and TLR are both required to disturb stem cell crosstalk and result in the disease for wildtype. TLR signalling alone was sufficient to perturb the crosstalk and drive disease progression for JAK mutated simulations. Furthermore, the model predicts the probability of disease onset for wildtype simulations that matches clinical data. These predictions might explain why patients who test negative for the JAK mutation can still be diagnosed with PMF, in which continual exposure to TPO and TLR receptor activation may trigger the initial inflammatory event that perturbs the bone marrow microenvironment and induce disease onset.

## Introduction

Primary myelofibrosis (PMF) is an idiopathic age-related clonal neoplastic disorder of haematopoiesis. Out of all the subsets of myeloproliferative neoplasms (including polycythaemia and essential thrombocytosis), PMF represents the highest morbidity and mortality and mean survival is estimated at less than 6 years post-prognosis (Bartalucci *et al*. 2013; Bartels *et al*. 2020).

More than 90% of myelofibrosis cases harbour somatic mutations in the driver genes JAK, CALR, or MPL that lead to constitutive activation of the JAK-STAT signalling pathways (Coltro *et al*. 2021). Following JAK activation, haematopoietic stem cells (HSCs) become hypersensitive to growth factors, rapidly proliferate and migrate from the bone marrow (BM) to sites of extramedullary haematopoiesis in the liver and spleen (Desterke *et al*. 2015; Hasselbalch. 2013). PMF is also accompanied by an event of chronic inflammation, which appears to be secondary considering that independent overexpression of each of the driver mutations in mice have been shown to recapitulate distinctive features of human disease mutations leading to JAK overactivity (Rumi et al., 2020).

Research has linked abnormally high levels of TGFβ, a potent inflammatory cytokine produced from atypical and dysmorphic megakaryocytes (MK) (Malara *et al*. 2018) in response to disrupted thrombopoietin (TPO) signalling in both patients and mouse models of the disease (Chagraoui *et al*. 2002; Melo-Cardenas *et al*. 2021). TPO, secreted from fibroblasts, is essential for MK differentiation and maintenance of HSCs (Cui *et al*. 2021) and signals through the JAK receptor (Varghese *et al*. 2017).

Current treatment protocols aim to disrupt the overactive STAT pathways (Stark and Darnell Jr. 2012; Pardanani *et al*. 2011) and diminish the effects of chronic inflammation, but are unable to influence the altered BM (Kuykendall *et al*. 2020). Within the BM microenvironment, HSCs reside in an endosteal niche and are engaged in constant crosstalk with neighbouring Mesenchymal Stem Cells (MSCs). MSCs secrete the chemokine CXCL12 and Vascular Cell Adhesion molecule-1 (VCAM1) which bind to their respective receptors, CXCR4 and VLA4, expressed upon the surface of HSCs. (Lubkova *et al*. 2011; Liekens *et al*. 2010). Chronic inflammation is believed to result in a break in any of these communication pathways which leads to HSC migration (Caocci *et al*. 2017).

ligand-activated stimulation of Toll-Like Receptors (TLRs) are well documented to induce chronic inflammation (Desterke *et al*. 2015; Fisher *et al*. 2017; Fisher *et al*. 2019; Fisher *et al*. 2021; Mascarenhas et al., 2022), leading to the hypothesis that unregulated TLR, and/or TPO signalling could be responsible for triggering chronic inflammation. This would then perturb the BM microenvironment and disrupt the crosstalk between HSCs and MSCs and lead to HSC migration and disease development. (Desterke *et al*. 2015). This hypothesis emphasises a major role of the microenvironment in the development of myeloproliferative neoplasms, and is interesting to consider in the context of patients who are “triple-negative” for one of the driver mutations (Tefferi and Vannucchi 2017), and for the 0.1 – 0.2% of the general population who carry the JAK mutation without showing overt signs of any myeloproliferative neoplasm(Nielsen *et al*. 2014).

It is well known that the presence of feedback circuits (or feedback loops) within biological systems ensures the maintenance of dynamical stability (Cornish-Bowden and Cárdenas. 2008; Qian and Beard. 2006). The idiopathic onset of PMF is believed to arise from an altered microenvironment that perturbs such feedback loops and results in disrupted communication between progenitor stem cells residing within the BM. The challenge of understanding disease onset therefore, requires a thorough understanding of how regulatory circuits control and regulate HSC-MSC crosstalk, and this challenge can be met using a computational systems approach. Construction of a computational model allows for a better understanding of the cause and effect of feedback loops and also provides a tool for making *in silico* predictions.

Quantitative and qualitative mathematical modelling formalisms have become powerful tools to study complex biological systems (Chaouiya *et al*. 2004; Montagud et al., 2019). Logical modelling presents a qualitative approach consisting of modelling regulatory networks using logical statements and allows for qualitative descriptions of dysregulated cellular signalling cascades resulting from altered microenvironments (Albert and Thakar. 2014; Naldi *et al*. 2015; Thieffry. 2007). Within this formalism, interesting insights between network structures and corresponding dynamical properties have been demonstrated (Remy and Ruet. 2008).

Enciso *et al*. (2016) published a logical model focusing on the inflammatory BM environment in response to TLR activation to capture NFKβ-dependent inflammation through disruption of the CXCL12/CXCR4 and VLA4/VCAM1 axis, accountable for the onset of Acute Lymphoblastic Leukaemia. We have elaborated upon this existing model to produce an extended model that captures the crosstalk and the key signalling pathways in the BM microenvironment whose alterations may lead to the onset of PMF.

## Results

### Logical modelling of the bone marrow microenvironment and stem cell crosstalk

A published logical model centred on the haematopoietic microenvironment between the HSC and MSC explaining lymphoblastic leukaemia (Enciso *et al*. 2016) was used as a starting model. This model, which contains 26 nodes and 80 interactions, encompasses the main communication pathways between HSPC and MSC within the BM, namely CXCL12/CXCR4 and VLA4/VCAM1 signalling. Whilst keeping with TLR signalling as an input, we replaced the original connexin43 input with TPO signalling due to its relevance in PMF. We also extended the HSC submodel by including JAK-STAT signalling pathway that is activated by either the TPO ligand or IL6 cytokine (Hasselbalch. 2013; Desterke *et al*. 2015; Fisher *et al*. 2019) which induces downstream PI3K/Akt signalling pathways (DelaRosa and Lombardo. 2010; Kawai and Akira. 2007; Bartalucci *et al*. 2013) and GCSF signalling (Schuettpelz *et al*. 2014; Brenner and Bruserud. 2019). We also removed the nodes CXCR7 and GFIL1 from the original HSC submodel as these nodes, despite being important in ALL, were considered to play a less important role in the development of PMF. Additionally, we have expanded the initial model by including an MK submodel, along with the main MK signalling pathways that are known to be dysregulated in PMF, that is with JAK-STAT signalling, TGFβ and PF4 production. Dysregulations in these pathways have been observed to alter the crosstalk between HSC and MSCs in the context of PMF.

We then set the output of the model as ‘PMF’ and included two pseudo output nodes, ‘HSC_proliferation’ and ‘HSC_migration’ that act as model readouts by concising characteristic cell fates relating to the disease.

A further defining feature of the model is that we have made use of multivalued nodes to discretise node activation states. Multivalued variables were associated with 9 nodes which allowed the capture of several threshold effects related to those nodes. The JAK, STAT, PI3K/Akt and NFKβ nodes of the HSC and MK submodels were given multivalued nodes. This is because 90% of PMF cases observe hyperactive JAK activity, resulting in the overactivation of downstream STAT, PI3K/Akt and NFKβ signalling within the HSC submodel (Rumi *et al*. 2006). We were able to capture these threshold effects within the model by allowing STAT, PI3K/Akt and NFKβ nodes to take their activity from the respective JAK activity, both in HSC and MK. The remaining node to be given a multivalued threshold was ‘HSC_proliferation’ which takes its value from HSC PI3K/Akt activity. HSC PI3K/Akt activity of 1 would activate ‘HSC_proliferation’ at its first, or basal level (considered as normal behaviour for stem cells); whilst HSC PI3K/Akt at its maximum threshold of 2 would then activate ‘HSC_proliferation’ at its highest level of 2. This second threshold reflects hyperproliferation as a consequence of overactive PI3K/Akt signalling, and this is associated with myeloproliferation (Skoda *et al*. 2015; Fisher *et al*. 2021) which is associated with PMF. In this way, we could set the output of the model as PMF, which is only activated when ‘HSC_proliferation’ has reached its maximum level of 2, and when ‘HSC_migration’ and ‘TGFβ_MK’ are activated.

The central axis within this expanded model is the CXCL12/CXCR4 chemokine pathway that now connects the MSC with HSC and MK submodels and plays essential roles in homeostasis and haematopoiesis (Moll and Ransohoff. 2010; Tamura *et al*. 2011). CXCR4 activation increases the affinity between VCAM1 expressed on the surface of MSC and its receptor VLA4, expressed by HSC and MK. Both pathways, CXCL12/CXCR4 and VCAM1/VLA4 are implicated with HSC migration (Lubkova *et al*. 2011; Moll and Ransohoff. 2010), and so the node ‘HSC_migration’ becomes activated when either the CXCL12/CXCR4 or VCAM1/VLA4 communication axis are broken. Otherwise, communication along these pathways triggers the PI3K/Akt and ERK signalling pathways that govern cellular activity and physiology (Yu and Cui. 2016).

Recent evidence links elevated secretion of pro-inflammatory cytokines within the BM niche in patients with PMF, including IL1 and IL6 (Desterke *et al*. 2015; Hasselbalch. 2013; Skoda *et al*. 2015), PF4(Gleitz *et al*. 2020; Melo-Cardenas *et al*. 2021) and TGFβ (Agarwal *et al*. 2016; Yao *et al*. 2019). IL1 provides an amplification role with chronic inflammation by activating PI3K/Akt and then NFKβ signalling, resulting in elevated production of both IL1, and ROS with the latter being counterbalanced by activation of FOXO transcription factors (Naka *et al*. 2008; Zhang *et al*. 2016). At the mesenchymal counterpart, TGFβ being produced by the MK has been reported to inhibit both VCAM1 (Park *et al*. 2000) and CXCL12 (Chagraoui *et al*. 2002), yet upregulates ERK signalling (Xue *et al*. 2020) and these interactions were also captured within the model. The inclusion of GSK3β and β-catenin in both HSC and MSC sub-models is relevant due to their roles as intermediates of signalling transduction and regulation of the main intracellular communication elements proposed in our network reconstruction (Figure 1).

**Figure 1.**
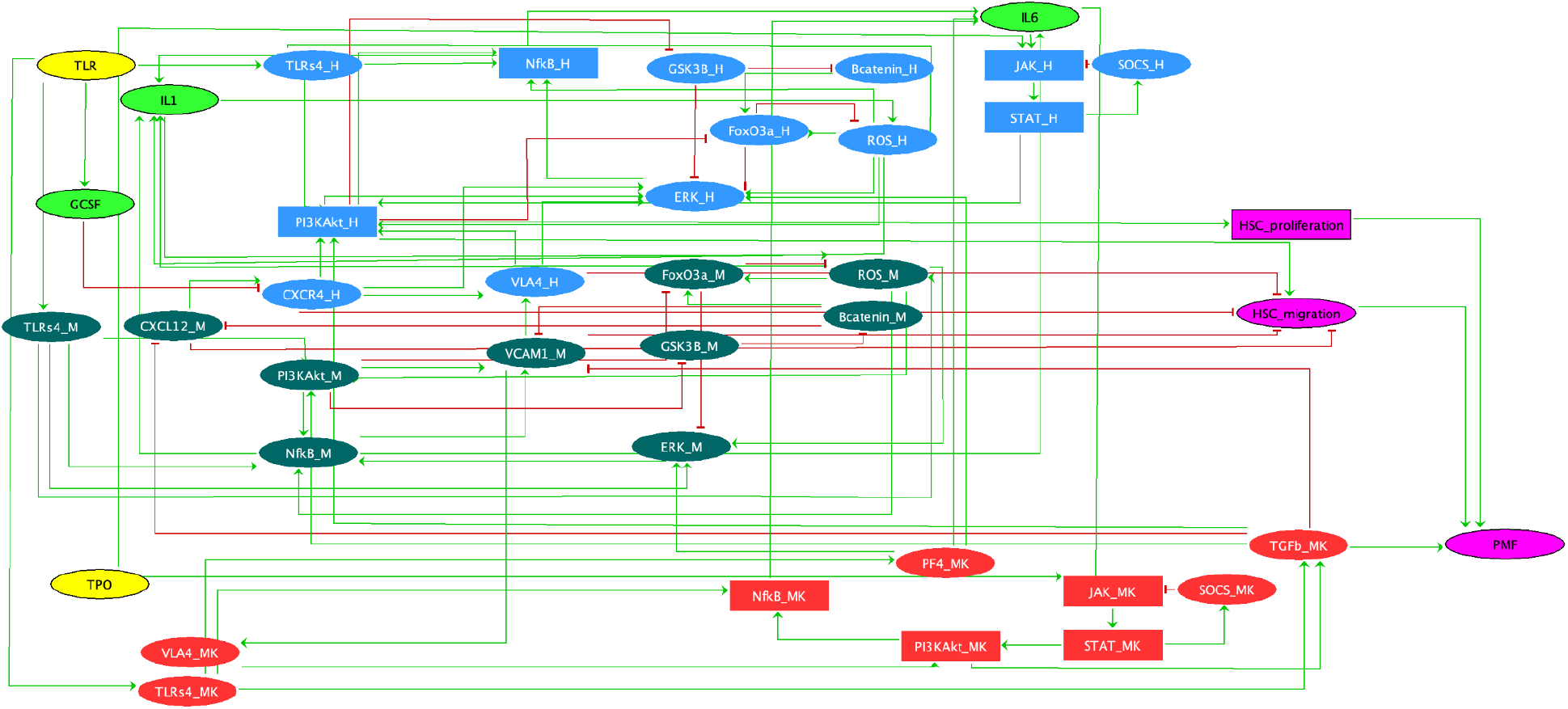
Regulatory graph capturing the crosstalk between HSC (blue nodes), MSC (dark green nodes), MK (red nodes) and inflammatory cytokines (light green nodes) belonging to the BM microenvironment. Nodes are connected by directed edges representing activations (green) and inhibitions (red). Regardless of sub-model specificity, ellipse-shaped nodes represent boolean nodes whilst rectangle-shaped nodes represent multivalue nodes. All logical rules and associated references can be found in Supplementary Table S1.

Modifying and expanding the model (Enciso *et al*. 2016) in order to account for myelofibrotic conditioning of the BM microenvironment (based on evidence in the literature, see Methods), we have obtained an extended model containing 40 nodes and 104 interactions (**Figure 1**).

This model was converted to a dynamical model by associating distinct logical rules to each node specifying the effect of the combination of the incoming interactions, based on biological knowledge. All logical rules and associated references can be found in **Supplementary materials Table S1**.

The model is available in XML format (GINML) on the GINsim Repository (http://ginsim.org/node/253).

The topology of the regulatory graph consists of a unique strongly connected component containing seven positive, and five negative functional circuits (see **Supplementary Table S2**, and **Supplementary Figure S3**). Some of these functional circuits interconnect all the cell submodels (HSC, MSC and MK), through the inflammatory cytokines submodel which implies an important role of inflammatory cytokines associated with disease onset. This emphasises a driving dynamics created by the cellular crosstalk between the several constituents of the BM microenvironment that is altered during the onset of PMF.

### Wildtype Attractor Analysis predicts TLR and TPO stimulation drive disease onset

We first investigated the attractors that describe the asymptotic behaviour of the model for wildtype simulations (without any mutation). For each combination of inputs, simulations resulted in a unique cyclic attractor. One cyclic attractor reveals a healthy phenotype in which PMF is averted (inputs [TPO, TLR] = [0, 0]). Another predicts a phenotype where the activity of the ‘PMF’ output oscillates between an active and inactive state (inputs [TPO, TLR] = [1, 1]), which reflects potential disease onset. Intermediate phenotypes are also observed in which the multivalued ‘HSC_proliferation’ and ‘HSC_migration’ nodes oscillate between inactive and active states, yet the remaining outputs are inactive (inputs [TPO, TLR] = [0, 1]). Finally, a phenotype involving the oscillation of the ‘HSC_proliferation’ node through all its states (0, 1 and 2), is observed and yet ‘HSC_migration’ and ‘PMF’ are otherwise inactive (inputs [TPO, TLR] = [1, 0]), and so PMF is averted (**Figure 2**).

**Figure 2.**
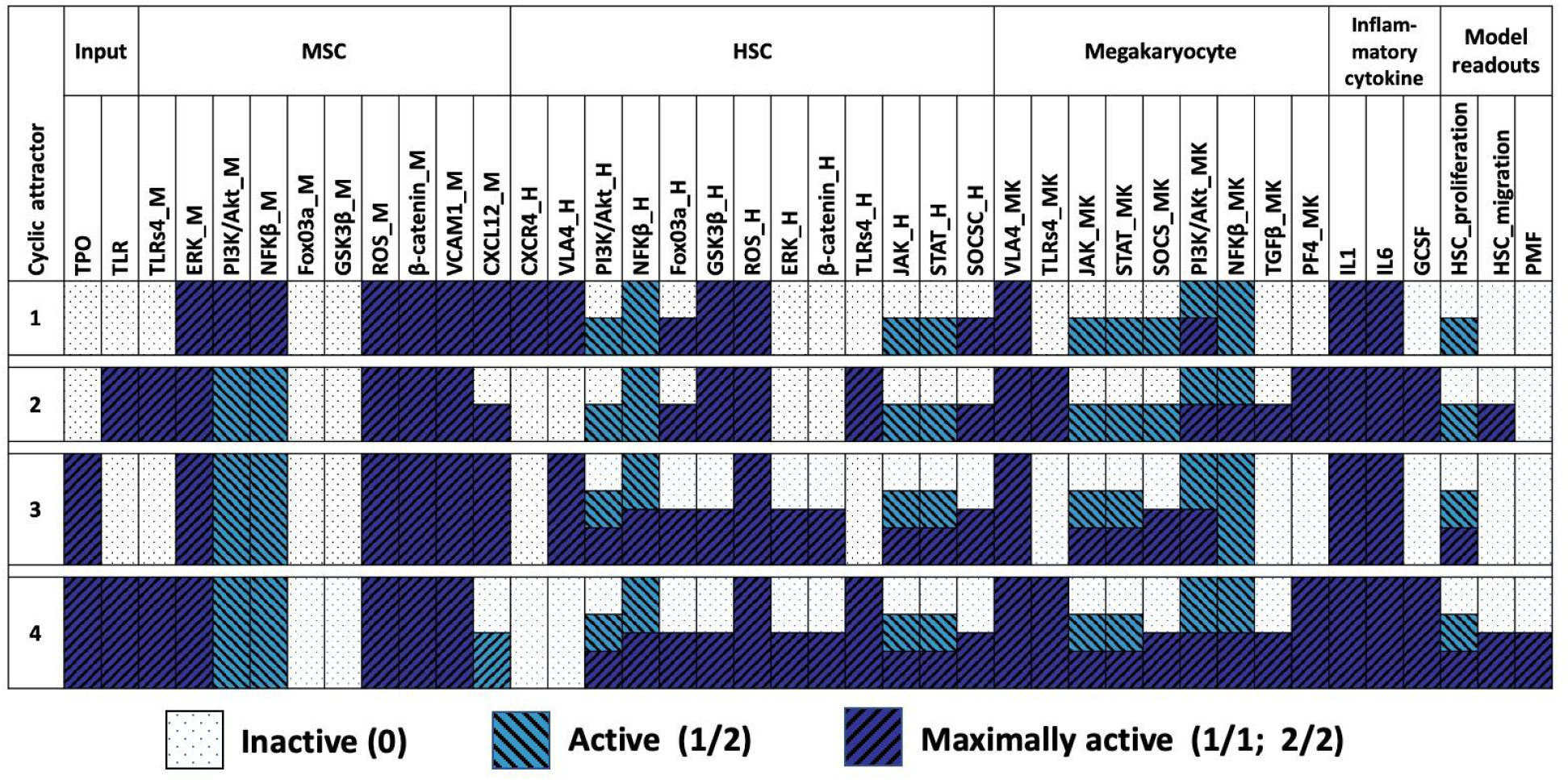
Description of the four attractors of the model (one for each input combination) in WT simulations. Each line is an attractor, and for each component (columns) a colour represents its state (see legend). Presence of several colours represent values of oscillating nodes. For example, in the first attractor with all inputs absent (first line), node ‘PI3K/Akt_H’ oscillates between 0 and 1, and ‘PI3K/Akt_MK’ is between 1 and 2.

In the absence of both inputs, MSC and HSC submodels remain connected via activation of CXCL12/CXCR4, and VCAM1 binding with VLA4 yet the multivalue ‘HSC_proliferation’ node oscillates between inactive and a state of partial activation but is never fully activated to its maximum threshold of 2.

The oscillatory range of ‘HSC_proliferation’ increases with the addition of either, or both inputs. When TPO is introduced as an input it binds to its receptor and combined with IL6, over-activates JAK and STAT signalling within the HSC and MK submodels. These oscillations drive both JAK receptors to now oscillate between the values of 0, 1 and 2, which then allows for a greater oscillatory nature of STAT and PI3K/Akt signalling which activates ‘HSC_proliferation’.

The presence of the TLR as a sole input triggers the activation of several positive feedback circuits that impact the HSC, MSC, MK and BM microenvironment. One of these cycles involves the direct activation of NFKβ and PI3K/Akt nodes from within the HSC and MK submodels resulting in the activity of ‘HSC_migration’ to oscillate. This is because TLR stimulates GCSF production which contributes to the inhibition of CXCR4 expression. TLR signalling also activates feedback cycles within the MK submodel that involve PI3K/Akt activation. PI3K/Akt activated at its maximal level results in the production of TGFβ from the MK which inhibits CXCL12 expression, yet upregulates PI3K/Akt signalling within the HSC which activates ‘HSC_migration’. When ‘PI3K/Akt_MK’ is inactive or at its basal level of 1, TGFβ is inactive and so CXCL12 remains activated and allows communication with the MSC via CXCR4.

When both inputs are introduced we observe oscillations of the ‘PMF’ output and so this attractor may describe a subset of PMF patients who don’t exhibit any driver mutation which may agree with the clinical data (Agarwal *et al*. 2016). In addition, TPO and TLR activate negative feedback circuits which result in the oscillatory activation of the ‘HSC_migration’ node. This cyclic attractor represents the communication breakdown of the HSC with the MSC and its migration out of the BM, as a result of TGFβ production. It is only in situations where the oscillatory activity of ‘HSC_proliferation’ is at its maximal level, and when the HSC is migrating, that we see the activation of the output node ‘PMF’. On the contrary, the onset of PMF is predicted to be avoided when the oscillatory activity of TGFβ remains inactive and when the HSC PI3K/Akt pathway remains in a status that is below its maximal activity. In order to get a finer description of the attractors, we ran stochastic simulations using MaBoSS (see methods). In particular, we wanted to calculate the probability of the ‘PMF’ output node being activated within the cyclic attractor obtained with both inputs present. As MaBoSS considers only Boolean systems, the multivalue node HSC_proliferation has been duplicated in two Boolean nodes, ‘HSC_proliferation_b1’ (centred on the threshold 1) and ‘HSC_proliferation_b2’ (centred on the threshold 2). When [TPO, TLR] = [1, 1], we observe that the nodes ‘HSC_migration’ and ‘HSC_proliferation’ follow similar trajectories and both have a 55-60% probability of being activated in this cyclic attractor. The probability of ‘HSC_proliferation_b2’ node (‘HSC_proliferation’ at its maximum level of 2) being activated in these conditions was predicted to happen with 17% probability, whilst activation of the ‘PMF’ output node was predicted to be activated with a probability of 8% which agrees with clinical data that 7-10% of PMF patients test negative for any of the driver mutations that overactive JAK signalling (**Figure 3**).

**Figure 3.**
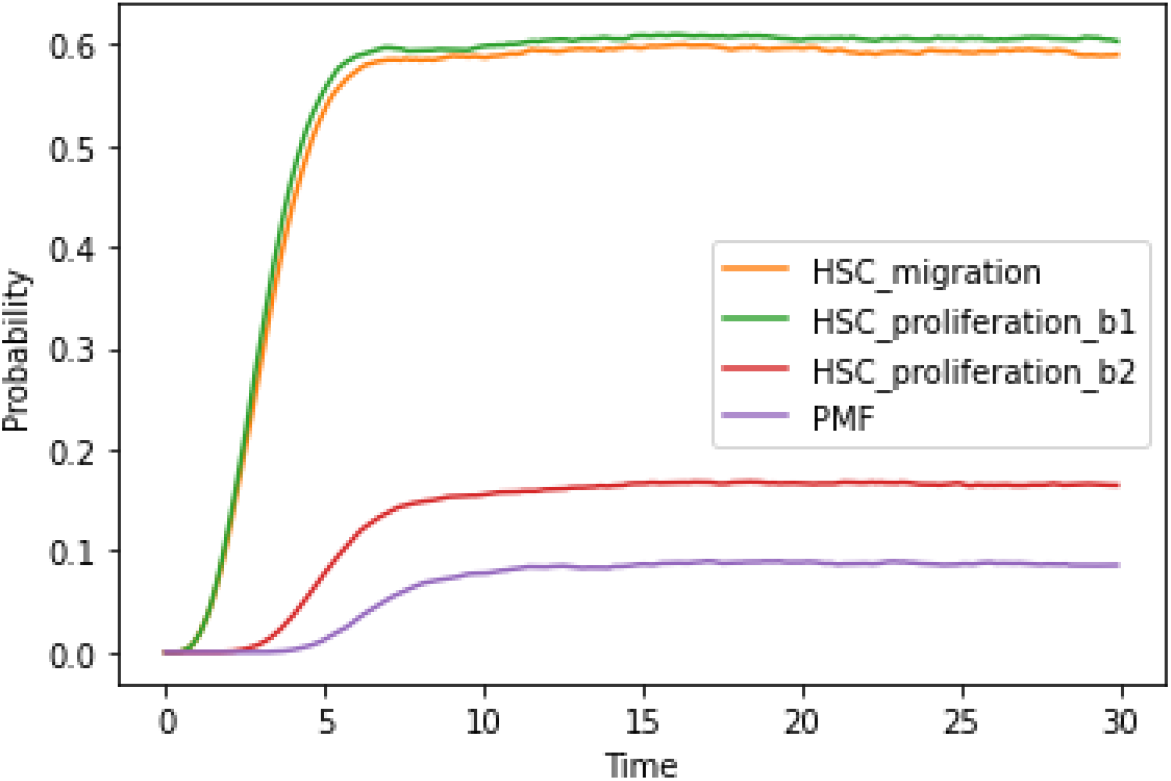
Probability of output nodes to be activated in the cyclic attractor from MaBoSS WT simulations, with both inputs present. ‘HSC_proliferation_b1’ represents the node has reached level 1 and ‘HSC_proliferation_b2’ is when the node has reached level 2.

We also investigated the probability of the oscillatory nodes JAK and TGFb being activated in the cyclic attractor (**Figure 5**). MaBoSS simulations showed an early increase in JAK activity, with the probability of being active at level 1 stabilising at 55% (after a transient peak at 70%). The probability of JAK achieving its level 2 activation is observed to be 30%. Once activated, ‘JAK_H’ remains activated because of sustained activation of IL6 which overrides the negative cycle involving SOCS inhibition of JAK. TGFβ release from MK was estimated to be 55% probable (**Figure 4**).

**Figure 4.**
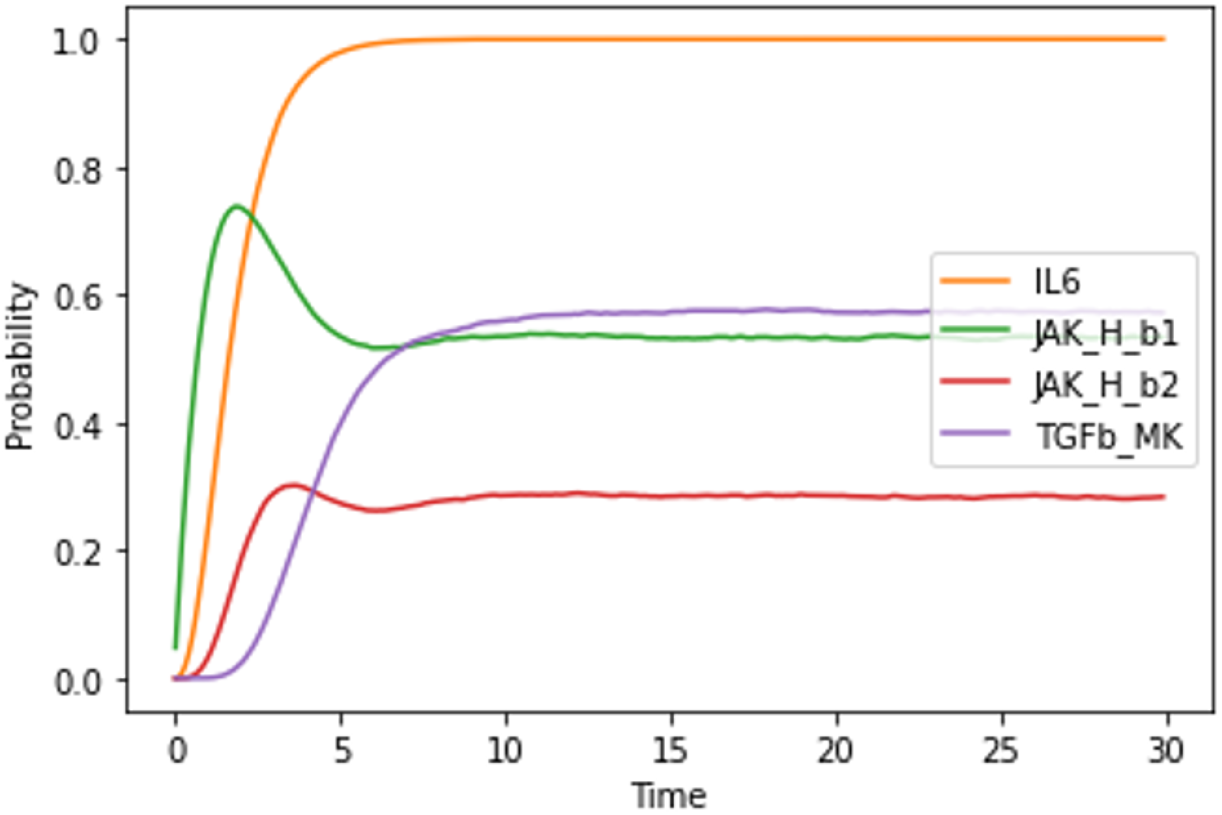
Probability of select nodes to be activated in the cyclic attractor from MaBoSS WT simulations, in the presence of both inputs.’JAK_H_b1’ represents the node reaching level 1 and ‘JAK_H_b2’ is when this multivalued node has reached level 2.

When comparing **Figures 3** and **4**, we observe a concomitant activation of HSC migration and proliferation (at level 1) with the progressive increase of IL6 activation and the probability of activating ‘JAK_H’ at its highest level is also increasing. This might explain the increase in probable activation of HSC overproliferation (‘HSC_proliferation_b2’). Meanwhile, the probability of TGFβ release increases, which explains the increasing probability of activating PMF.

To conclude, these simulations reveal that in the absence of the JAK mutation (WT situation), both inputs TPO and TLR have to be present for PMF to be activated, probably arising from a JAK overactivation which cascades inflammatory feedback circuits through IL6 and TGFβ pathways.Jak Mutated Analysis predicts the presence of TLR alone is sufficient to drive disease onset

All known driver mutations associated with PMF induce overactive JAK activity and downstream STAT signalling, and are observed in 90% of PMF cases, whilst 0.1 – 0.2% of the population harbour the JAK mutation but do not display signs of the disease (Agarwal *et al*. 2016). In an attempt to explain why some patients with the overactive JAK activity eventually manifest with PMF whilst others do not, we simulated ‘JAK_H’ and ‘JAK_MK’ KI mutants. In this scenario, the model displays 4 fixed points, one per combination of input nodes (**Figure 5**). These fixed points reveal two healthy phenotypes (inputs [TPO, TLR] = [0, 0]; [1, 0]), and two that activate PMF (inputs [TPO, TLR] = [0, 1]; [1, 1]).

**Figure 5.**
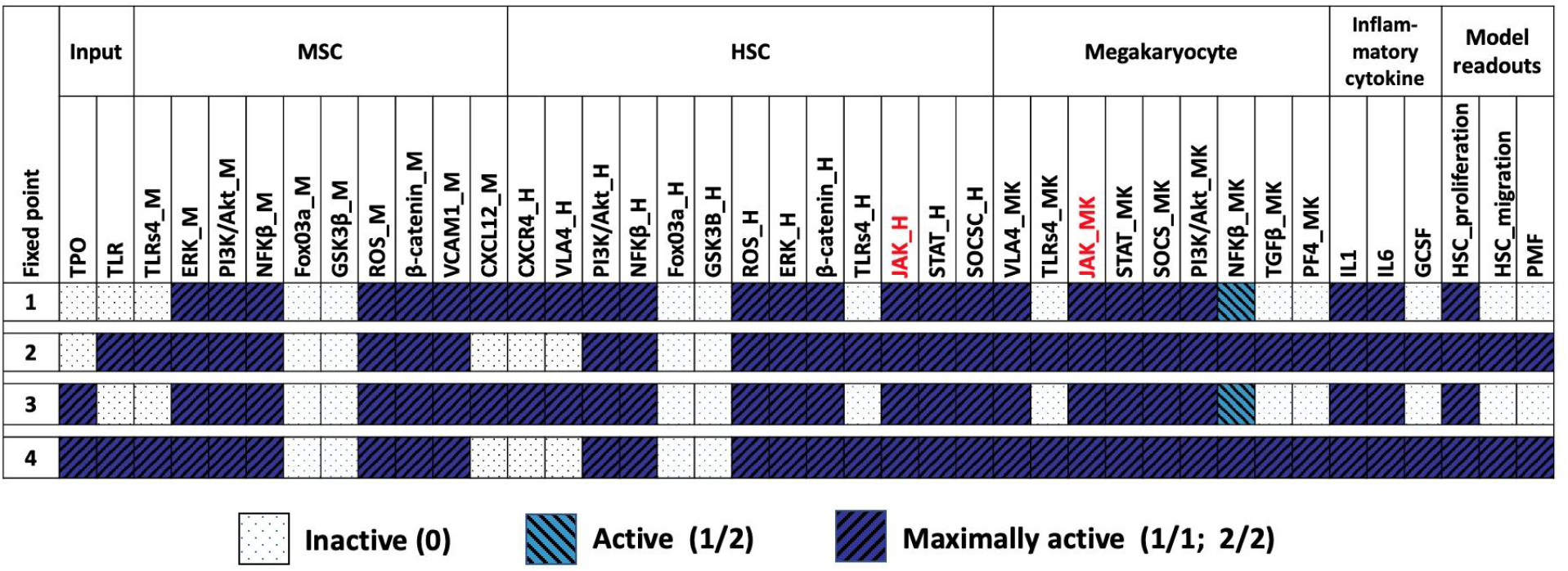
Fixed points for the JAK ectopic simulation (mutated nodes are marked with red text).

Simulations reveal that the JAK mutation activates positive feedback cycles in both the HSC and MK submodels that involve activation of STAT and downstream PI3K/Akt and NFKβ signalling which then activates IL6 which feeds forward to sustain the activation of the initial JAK receptor. Overactive ‘PI3K/Akt_H’ activity is responsible for HSC hyperproliferation, as this multivalued node has reached its maximum level. The HSC is not predicted to migrate because of a sustained connection with the MSC submodel following the activation of mesenchymal PI3K/Akt which promotes the activation of VCAM1 and allows the reception of VLA4. Furthermore, TGFβ and GCSF are inactive and so CXCL12/CXCR4 remain active in the absence of TLR. This could explain why the subset of the population who exhibit any driver mutation that overactives the JAK receptor avoids the onset of PMF.

Following activation of both inputs for JAK mutated simulations, TLR signalling augments the JAK burden in the MK submodel causing overactivation of PI3K/Akt signalling and this time, NFKβ signalling. TPO also augments JAK stimulation, and TLR activates GCSF and PF4, and inactivates CXCR4 and VLA4, resulting in the loss of communication between the HSC and MSC submodels. Hyperactive JAK signalling within the HSC submodel led to overactive PI3K/AKt, and when combined with the activation of ‘HSC_migration’ and TGFβ, led to the activation of the output node ‘PMF’, at a calculated probability of 100% (**Supplementary materials Figure S4**).

Again, these results agree with the literature that shows activation of TPO and JAK are both required to induce a myelofibrotic phenotype (Besancenot *et al*. 2014). Furthermore, PF4 (Gleitz *et al*. 2020) and TGFβ (Chagraoui *et al*. 2002; Zingariello *et al*. 2013; Agarwal *et al*. 2016) are critically implicated with disease progression. In conclusion, JAK (KI) mutated simulations predict that TLR signalling is sufficient to perturb the microenvironment and disrupt HSC-MSC crosstalk to drive disease onset.

## Discussion and Conclusion

Primary myelofibrosis is an untreatable age-related disorder of haematopoiesis in which a break in the crosstalk between progenitor HSCs and neighbouring mesenchymal stem cells (MSC) caused HSCs to rapidly proliferate and migrate out of the bone marrow. In this work, we have elaborated and adapted a pre-existing logical model of lymphoblastic leukaemia (Encisco et al., 2016) to account for the BM microenvironment with the primary aim of capturing stem cell cross-talk between HSC, MSC and megakaryocyte progenitors. We then used the model to capture the dysregulation of the cell crosstalk following perturbation of the BM microenvironment due to stimulation with TPO, and/or activation of TLR, for wildtype and in ectopically JAK mutated situations. We chose to include TPO and TLR signalling as inputs due to their roles in mediating chronic inflammation.

The model predicted that for wildtype scenarios, the presence of both inputs (TPO and TLR) resulted in a probability of disease. TLR signalling was predicted to activate PF4, which agrees with (Gleitz *et al*. 2020) who report that elevated expression of PF4 links inflammation with progression of BM fibrosis. The authors also report that the absence of PF4 reduces the extent of BM remodelling in presence of overactive JAK pathways, which our model has qualitatively captured (**Figure 2; Figure 5**). Our results suggest that TLR signalling cascades positive and negative feedback circuits that are responsible for triggering an initial event of chronic inflammation through the activation of PF4 and TGFβ. TPO signalling cascaded further positive and negative feedback circuits that facilitate HSC hyperproliferation, independent of the JAK mutation. These predictions might explain why patients who test negative for the JAK mutation can still be diagnosed with PMF, in which continual exposure to TPO and TLR receptor activation may trigger the onset of PMF. JAK mutated (KI) simulations revealed that TLR alone was sufficient to drive myelofibrosis. The model also predicted that in the presence of JAK mutation and absence of TLR, PMF was averted which supports the observation that 0.1 – 0.2% of the general population harbour the JAK mutation without showing overt signs of any myeloproliferative disease (**Figure 5**). We hypothesise that the probability of TLR and TPO signalling increases with time, explaining why the onset of PMF is age-related.

Mathematical modelling aims at abstracting and understanding the effects of biological perturbations to suggest ways to intervene and reestablish homeostasis. This is challenging to achieve within any modelling framework. The model presented in this work contains limited, but sufficient biological interactions that play a role in the onset of PMF. We sought to capture as much information as possible while maintaining enough abstraction to obtain a manageable model of reasonable size for the analysis to be performed. This has allowed the construction of a model that agrees qualitatively -and in some respects even quantitatively-with experimental and clinical data. As a limitation, we are modelling crosstalk between single cells, while in reality, multiple cells reside within a niche which our approach does not take into account. It is therefore essential to explore the dynamics at the scale of interacting cell populations, taking into account their response, death and interactions. This requires a proper computational description of heterogeneous interacting cell populations, and this model could be the basis of such modelling at the cell population level microenvironment.

In conclusion, we have built an intra- and intercellular logical model that is able to capture the crosstalk between distinct progenitor stem cells within the BM microenvironment. The model is able to predict and explain the onset of myelofibrosis for JAK-positive and JAK-negative cases. More generally, this model could be useful for the study of other haematopoietic diseases.

## Methods

### Logical Modelling

Logical modelling is a qualitative formalism developed to study the dynamics of complex biological networks (Albert and Thakar. 2014; Abou-Jaoudé *et al*. 2016). Biological information is extracted from the literature and/or experiments and abstracted into a regulatory graph (RG), a directed signed graph whose nodes represent selected biological species, connected by regulatory edges (positive edges for activations, negative for inhibitions). Input nodes (nodes with only outgoing interactions) usually represent the environment of the cell or any extracellular signal, and output nodes (that only have incoming interactions and thus no impact on the model) usually serve as a readout of the model. Outputs can reflect an observed phenotype such as ‘proliferation’ and/or ‘migration’, as examples.

To introduce the dynamics, each node is associated with a discrete variable, denoting its state (in the binary case, 0 (inactive) or 1 (active)). Nodes can also be assigned with multivalued values that precise different thresholds, or activation levels. The global state of the network is represented by a vector of *n* elements, where *n* represents the number of nodes in the network. Each node is associated with logical rules that describe the target value of the variable with respect to the state of the regulators of the node. These logical rules set the dynamics and the asynchronous updating strategy provides non-deterministic dynamics by updating a single component at each step (see example; Figure 7). Trajectories ultimately end within an attractor, i.e. terminal strongly connected component of the graph and represent the asymptotic behaviours of the system. They can either be fixed points (attractors of size one), or cyclical attractors.

Simulations of mutants consist in modifying logical rules. Knock-out (KO) mutants are simulated by blocking the logical rule of the desired nide to 0, and Knock-in (KI) mutants by fixing it to its maximal level or active at the desired threshold for multivalued nodes (overexpression).

**Figure 7.**
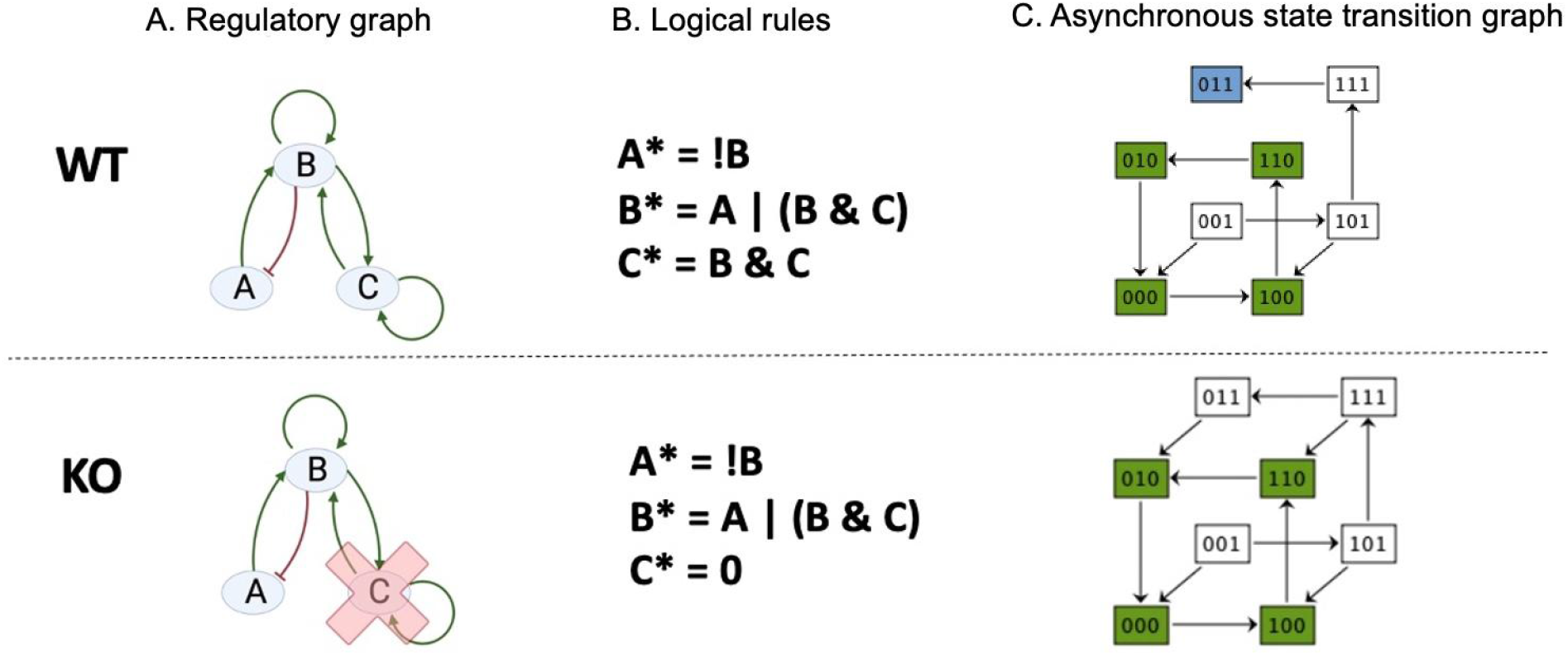
Toy model illustrating the logical modelling methodology. The first row depicts a wildtype (WT) situation, and the bottom row depicts a KO mutation of node C. **A**) The regulatory graph (RG) consists of 3 nodes, A, B and C, and 6 edges, 5 activatory edges (green arrows) and one inhibitory edge (the red arrow). **B**) Logical rules (*indicates possible future active state when the following logical rules are met). When node C has been knocked out (mutant version), its rule is blocked to 0. **C**) Asynchronous State Transition Graphs (STG). The asynchronous STG contains two attractors, a single fixed point [ABC] = [011] (coloured blue) and a cyclic attractor composed of the states {[000]; [100]; [110]; [010]} (coloured green). Asynchronous dynamics of the mutant C KO loses the fixed point and only results in the same cyclic attractor (coloured green).

The logical model was built using GINsim software and is available in XML format (GINML) on the GINsim Repository (http://ginsim.org/node/253, also provided in Supplementary File 1). All attractors, belonging to either the wildtype and mutant simulations (double mutation JAK_H KI and JAK_MK KI)were sought and characterised using GINsim in conjunction with the CoLoMoTo toolbox (Levy *et al*. 2018) using Python.

### Feedback Circuit Analysis

Feedback circuits are known to play a major role in the control of cell signalling dynamics. For instance, the Thomas’ rules state that the presence of positive (resp. negative) circuits are necessary prerequisites to generate multistability (resp. sustained oscillations) (Thomas. 1981; Thieffry. 2007).

When embedded withina RG, a circuit is regulated by some external nodes and thereby its activity is affected. Circuit functionality contexts indicate the part of the space in which the positive (resp. negative) circuit is able to generate local multistability (resp. oscillations), and we say that the circuit is functional. These contexts are defined in terms of constraints on the values of external nodes. The “Analyse circuits” tool of GINsim lists all the functional circuits present in a logical model and their context of functionality.

### MaBoSS Simulations

MaBoSS (Markovian Boolean Stochastic Simulator; Stoll *et al*. 2017) provides an environment for simulating continuous/discrete time Markov processes based on logical networks by applying the Gillespie algorithm. MaBoSS calculates the time evolution of the probability of node states being activated and so was used to see if the model could predict the probable onset of PMF with, and without JAK mutation as a means of model validation. In addition, global and semi-global characterizations of the whole system are computed. Simulation parameters involve running 50,000 simulations performed to compute statistics and the maximum time representing the duration of the trajectory was set to 30. For wildtype (JAK KO) simulations, the initial states of all nodes were set to zero, apart from the inputs; which were both set to be 100% active. The inputs for the JAK mutated simulations account for TLR which was 100% probable of being activated. The initial states of non-input nodes were then set to inactive apart from ‘JAK_MK’ and ‘JAK_HSC’, as these were ectopically mutated to their maximum levels. The average temporal evolution of selected nodes of the model were then plotted.

## Declarations

### Ethics approval and consent to participate

Not applicable

### Consent for publication

Not applicable

### Data availability

The data underlying this article are available within the GINsim repository at

http://ginsim.org/node/253

### Authors contributions

ED thought up and designed this work. ER supervised the modelling methodology. SC built, ran and analysed the model and wrote the manuscript. All authors read and approved the final manuscript.

## Acknowledgements

The authors would like to thank A. Naldi for his assistance with GINsim, P. Monteiro for uploading the model to the GINsim model repository, F. Zaccagnino for assisting with MaBoSS and B. Habermann for proofreading the manuscript.

