## Supplementary materials for "Logical modelling of myelofibrotic microenvironment predicts dysregulated progenitor stem cell crosstalk"

**Supplementary Table S1: Logical rules**

| Nodes | Values | Logical rules | References |
|---|---|---|---|
| **TPO** | 0/1 | Input | de Graaf & Metcalf (2011) |
| **TLR** | 0/1 | Input | Kawai & Akira (2007) |
| **TLRs4_M** | 1 | TLR | English & Mahon (2011) |
| **ERK_M** | 1 | !FoxO3a_M & (ROS_M \| TLRs4_M \| PF4) | Pan et al., (2017)<br>Zhang et al., (2017)<br>Banerjee et al., (2006)<br>Ryu et al., (2010) |
| **PI3KAKt_M** | 1 | TLRs4_M \| ROS_M \| TGFb_MK | DelaRosa & Lombardo (2010) |
| **NFKβ_M** | 1 | ROS_M \| PI3KAkt_M \| ERK_M \| TLRs4_M | Bjørn & Hasselbach (2015)<br>Desterke et al., (2015) |
| **FoxO3a_M** | 1 | (Bcatenin_M \| ROS_M) & !PI3KAkt_M | Ghaffari (2008);<br>Hay (2011) |
| **GSK3B_M** | 1 | !PI3KAkt_M | Hermida et al., (2008) |
| **ROS_M** | 1 | IL1 \| TLRs4_M \| !FoxO3a_M | Dinarello (2011) ;<br>Naka et al., (2008) |
| **β-catenin_M** | 1 | !GSK3B_M | Tamura, Sato & Nashimoto, M. (2011) |
| **VCAM1_M** | 1 | !Bcatenin_M \| NfkB_M \| PI3KAkt_M \| !TGFb_MK | Malhotra & Kincade (2009);<br>Hu et al., (2009);<br>Park et al., (1995) |

| | | | |
|---|---|---|---|
| **VLA4_H** | 1 | VCAM1_M \| CXCR4_H | Moll & Ransohoff (2010). Lubkova et al., (2011) |
| **CXCL12_M** | 1 | !TGFb_MK \| !Bcatenin_M | Yao et al., (2019); Jung et al., (2006); Tamaro et al., (2011) |
| **CXCR4_H** | 1 | CXCL12_M & !GCSF | Le Bousse-Kerdilès (2012); Xu et al., (2005) Greenbaum & Link (2011) |
| **PI3KAKt_H** | 1 | (CXCR4_H \| ROS_H \| TLRs4_H \| VLA4_H \| TGFb_MK) & STAT_H:1 | Troutman et al., (2012); (Kato et al., 2009); Bartalucci et al., (2103) |
| | 2 | (CXCR4_H \| ROS_H \| TLRs4_H \| VLA4_H \| TGFb_MK) & STAT_H:2 | Quintás-Cardama et al., (2013); Zhan et al., (2020) |
| **NFKb_H** | 1 | ROS_H & !PI3KAkt_H:2) \| (ERK_H & !PI3KAkt_H:2) \| (TLRs4_H & !PI3KAkt_H:2) | Bjørn & Hasselbach (2015); Desterke et al., (2015) |
| | 2 | PI3KAkt_H:2 | Zimran et al., (2018) Hasselbalch (2013) |
| **FoxO3a_H** | 1 | (Bcatenin_H \| ROS_H) & !PI3KAkt_H | Ghaffari (2008) |
| **GSK3B_H** | 1 | !PI3KAkt_H:2 | Falconi et al., 2016) |
| **ROS_H** | 1 | !FoxO3a_H \| IL1 \| TLRs4_H | Kawai & Akira (2007) |
| **ERK_H** | 1 | ((CXCR4_H & PI3KAkt_H) \| ROS_H \| VLA4_H \| PF4) & !(FoxO3a_H \| GSK3B_H) | Stivala et al., (2019) |
| **bcatenin_H** | 1 | !GSK3B_H | Tamura, Sato & Nashimoto, M. (2011) |
| **TLRs4_H** | 1 | lTLR | Kawai, T., & Akira, S. (2007) |

| | | | |
|---|---|---|---|
| **JAK_H** | 1 | (IL6 & !TPO & !SOCS_H) \| (TPO & !IL6 & !SOCS_H) \| (!IL6 & !TPO & !SOCS_H) | Fisher et al (2019)<br>Malara et al., (2018) |
| | 2 | TPO & IL6 & !SOCS_H | Hasselbalch (2013)<br>Johnson et al., (2018) |
| **STAT_H** | 1 | JAK_H:1 | Stark & Darnell (2012) |
| | 2 | JAK_H:2 | Stark & Darnell (2012) |
| **SOCS_H** | 1 | STAT_H | Stark & Darnell (2012) |
| **VLA4_MK** | 1 | VCAM_1 | Moore (2002) |
| **TLRs4_MK** | 1 | lTLR | Kawai, T., & Akira, S. (2007) |
| **JAK_MK** | 1 | (IL6 & !TPO & !SOCS_MK) \| (TPO & !IL6 & !SOCS_MK) \| (!IL6 & !TPO & !SOCS_MK) | Fisher et al., (2019);<br>Malara et al., (2018) ;<br>Johnson et al., (2018) |
| | 2 | TPO & IL6 & !SOCS_MK | Hasselbalch (2013) |
| **STAT_MK** | 1 | JAK_MK :1 | Stark & Darnell (2012) |
| | 2 | JAK_MK:2 | Stark & Darnell (2012) |
| **SOCS_MK** | 1 | STAT_MK | Stark & Darnell (2012) |
| **PI3KAKt_MK** | 1 | (VLA4_MK & !STAT_MK) \| (STAT_MK:1 & !VLA4_MK) | Troutman et al., (2012);<br>Kato et al., (2009);<br>Bartalucci et al., (2013) |
| | 2 | STAT_MK:2 \| (VLA4_MK & !STAT_MK) \| (STAT_MK:1 & !VLA4_MK) | Quintás-Cardama et al., (2013);<br>Zhan et al., (2020) |
| **NFKb_MK** | 1 | (TLRs4_MK & !PI3KAkt_MK) \| (PI3KAkt_MK & !TLRs4_MK) | Bjørn & Hasselbach (2015);<br>Desterke et al., (2015) |

| | | | |
|---|---|---|---|
| | 2 | TLRs4_MK & PI3KAkt_MK | Zimran et al., (2018)<br>Hasselbalch (2013) |
| **TGFb** | 1 | PI3KAkt_MK:2 & TLRs4_MK | Malara *et al.* 2018 |
| **PF4** | 1 | TLRs4_MK | Cognasse et al., (2015) |
| **IL1** | 1 | ROS_H \| ROS_M \| NfkB_M \| NfkB_H | Hasselbalch, H. C. (2013) |
| **IL6** | 1 | NfkB_M \| NfkB_MK \| PF4 | Hasselbalch, H. C. (2013) |
| **GCSF** | 1 | TLR | Schettpelz et al., (2014) |
| **HSC_migration** | 1 | (PI3KAkt_H & (!CXCL12_M \| !CXCR4_H)) \| (PI3KAkt_H & (!VCAM1_M \| !VLA4_H)) | Moll & Ransohoff (2010)<br>Lataillade et al., (2008) |
| **HSC_proliferation** | 1 | PI3KAkt_H:1 | Jason & Cui (2016) |
| | 2 | PI3KAkt_H:2 | Bartalucci et al., (2013) |
| **PMF** | 1 | HSC_migration & TGFb_MK & HSC_proliferation:2 | Malara et al., (2018);<br>Lataillade et al., (2008) ;<br>Bartalucci et al., (2013) |

The model variables (column 1) can take values 0, 1 or 2 (for the multi-valued variables: PI3KAKt_H' NFKβ_H, JAK_H, STAT_H, PI3KAKt_MK, NFKβ_MK, JAK_MK, STAT_MK and HSC_proliferation; column 2). Logical rules are associated to each variable, (column 3), justified from the literature (column 4).

**Supplementary File 1:** logical model in ginml format

**Supplementary Table S2: Functional circuits present within the regulatory network**

| Circuit | Nodes contained within circuit | Sign |
|---|---|---|
| **1** | IL1 - ROS_M | positive |

| | | |
|---|---|---|
| **2** | PI3KAKt_H – FoxO3a_H – ROS_H | positive |
| **3** | IL6 – JAK_H[1] – STAT_H[1] – PI3KAKt_H[1] – NFKβ_H (1/8) | positive |
| **4** | PI3KAKt_MK[1] – TGFβ_MK – PI3KAKt_H[1] – NFKβ_H – IL6 – JAK_MK[1] – PI3KAKt_M | positive |
| **5** | NFKβ_M – IL6 – JAK_MK[1] – STAT_MK[1] – PI3K/AKt_MK[1] – TGFβ_MK – PI3KAKt_M | positive |
| **6** | ROS_M – PI3KAKt_M – FoxO3A_M | positive |
| **7** | IL1 – ROS_H | positive |
| **8** | ROS_M - FoxO3A_M | negative |
| **9** | JAK_MK – STAT_MK[1] – SOCS_MK (1/2) | negative |
| **10** | Fox03a_H – ROS_H | negative |
| **11** | JAK_H– STAT_H – SOCS_H (1/2) | negative |
| **12** | VCAM1_M – VLA4_MK – PI3KAKt_MK[1] – TGFb_MK (1/2) | negative |

Functional regulatory circuits present in the model. Nodes preceded by '_H', '_M' and '_MK' represent belonging to the HSC, MSC and MK sub model respectively; otherwise, belong to the BM microenvironment. Mutlivalue nodes can result in several circuits because the various activation statuses of such nodes propagate different edge thresholds and each circuit considers one threshold of each multilevel edge threshold. The number of circuits activated by such multivalued nodes are represented within parenthesis.

**Supplementary Figure S3: Topological structure of the Regulatory network at the cell scale**

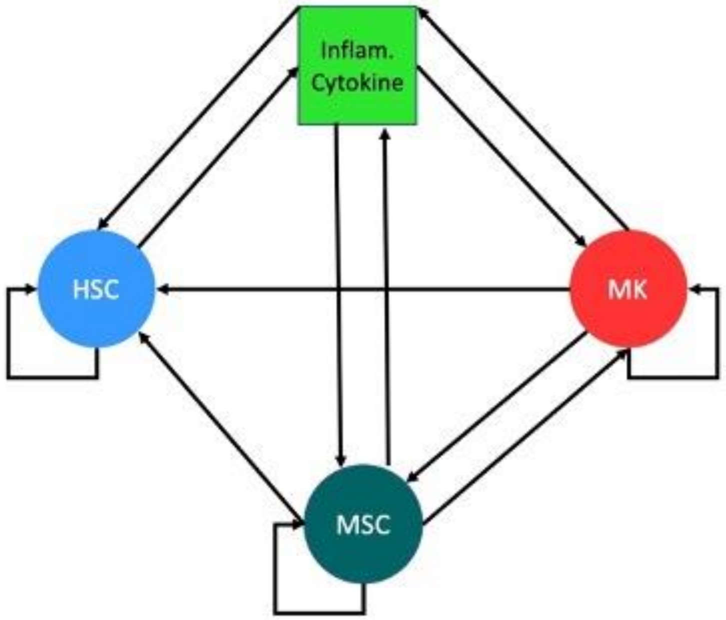


Representation at the scale of the cell of the RG. Nodes represent each submodel of the logical model encompassing HSc, MSC, MK and the complement of inflammatory cytokines within the bone marrow. Functional regulatory circuits that connect each submodel, listed in **supplementary materials S3** are represented by arcs.

**Supplementary Figure S4: MaBoSS simulation of ectopically JAK mutated model in the presence of both inputs**.

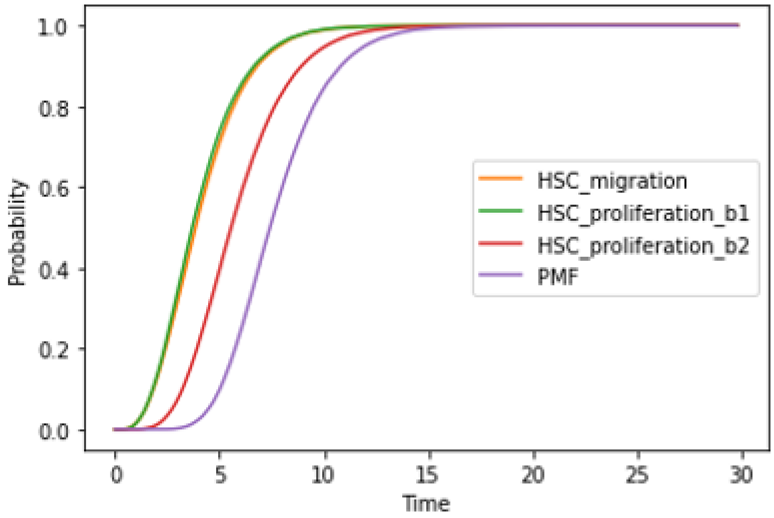

MaBoSS simulation of ectopically JAK mutated model in the presence of both inputs. HSC proliferation is a multivalued node and its thresholds are represented as ‘b1’ when its first threshold of ‘1’ has been reached, and ‘b2’ denotes when the node has reached its maximum threshold of ‘2’.
